# Improved RNA homology detection and alignment by automatic iterative search in an expanded database

**DOI:** 10.1101/2022.10.03.510702

**Authors:** Jaswinder Singh, Kuldip Paliwal, Jaspreet Singh, Thomas Litfin, Yaoqi Zhou

## Abstract

Unlike 20-letter-coded proteins, RNA homologous sequences are notoriously difficult to detect because their 4-letter-coded sequences can quickly lose their sequence identity. As a result, employing secondary structures has been found necessary to improve the sensitivity and the accuracy of homolog search. However, exact secondary structures often are not known. As a result, Rfam, the de facto gold-standard of RNA homologous families, has to rely on manual curation and experimental secondary structure if available. Here, we showed that using a combination of BLAST and iterative INFERNAL searches along with an expanded sequence database leads multiple sequence alignments (MSA) that are comparable to those provided by Rfam MSAs, according to secondary structure extracted from mutational coupling analysis and alignment accuracy when compared to structure alignment. The fully automatic tool (RNAcmap2) allows making homolog search, multiple sequence alignment, and mutational coupling analysis for any non-Rfam RNA sequences with Rfam-like performance.

## Introduction

Homology search is the first step for getting a clue about an RNA with an unknown function. A set of homologous sequences, if found, will allow not only functional inference but also provide structural consensus for improving secondary structure prediction, for example, as in RNAaliFold [1] and CaCoFold [2]. Multiple Sequence Alignment (MSA) of homologous sequences can further reveal the pairs of co-varying bases, indicating conserved secondary and tertiary structural features [3, 4, 5, 6, 7]. Sequence profiles and correlated mutations generated from multiple sequence alignment have been found useful for significantly improving prediction of RNA secondary structure [8] and solvent accessibility [9, 10, 11] by deep learning.

Existing homology search and alignment tools can be classified into two categories: sequence-based and profile-based. BLAST-N [12] is a popular homolog search tool based on the heuristic Smith-Waterman algorithm [13] for similarity search between two sequences. Profile-based methods include nhmmer [14] and INFERNAL [15]. nhmmer employs profile Hidden Markov Models (profile HMM) to search a query against a reference library, where the query can be a single sequence, MSA or profile HMM. INFERNAL incorporates secondary structure along with a query sequence/MSA and builds a covariance model (CM) equivalent to the sequence-based profile HMM. The CM is then utilized to search against a reference library to find even more remote homologs. In general, profile-based homolog searches are more sensitive in detecting remote homologs than sequence-based searches and including secondary structure in profile-based searches (INFERNAL) is more sensitive than using profile only (nhmmer). INFERNAL and manual curation were utilized to obtain Rfam homologous sequences along with their MSAs within an RNA family, by using experimentally validated or predicted structural information [16]. Rfam alignment has been considered as the “de facto” gold standard for MSA. However, the exact secondary structure of an RNA is often unknown, and manual curation is too slow to meet the demand from exponentially growing RNA sequences [17].

Recently, we developed a fully automated pipeline RNAcmap [18] to obtain an aligned set of homologs. RNAcmap used the BLAST-N and INFERNAL tools against the NCBI’s nucleotide database [17] for homology search. However, for sequences annotated in Rfam, the base pairs detected by mutational coupling from RNAcmap aligned sequences is not as accurate as those from Rfam aligned sequences.

This work improves the RNAcmap pipeline with one additional search and expanded reference database. The proposed pipeline (RNAcmap2) first search by BLAST-N followed by two additional INFERNAL searches. Moreover, we expand the NCBI’s nucleotide database by including NCBI’s metagenomics and patent sequences. RNAcmap2 yields the alignment and structural co-variation signals as accurate as Rfam supplied MSAs. Unlike Rfam, RNAcmap2 can produce MSA for any RNA sequences with > 50% improvement in F1-score over RNAcmap on 127 non-Rfam RNAs for predicted contact base pairs, according to direct coupling analysis [3, 4, 5, 6, 7].

## Materials and methods

### Datasets

For benchmarking, we downloaded all high-resolution (<3.5Å) X-ray structures that include RNA chains from Protein Data Bank [19] on October 3, 2020. Individual RNA chains were extracted from RNA-RNA and RNA-protein complex structures using a PDBParser from Biopython [20]. To remove the redundancy within RNA chains, we used CD-HIT-EST [21] at the lowest allowed sequence identity cut-off of 0.8. This led to a total of 245 non-redundant and high-resolution RNA chains with minimum and maximum sequence length of 33 and 418 respectively. RNA secondary structure labels for these chains were obtained from their 3D structural files using DSSR [22].

The method performance of structure-prediction tools strongly depends on the number of effective homologous sequences N*_eff_*. Thus, we divided our dataset into four categories: No-hit, Low N_*eff*_ (1 ≤ N*_eff_* < 10), Medium N*_eff_* (10 ≤ N_*eff*_ < 50), and High N*_eff_* (50 ≤N*_eff_*) RNAs as shown in Table 1, where the N*_eff_*-value was obtained from RNAcmap [18] with the NCBI nucleotide database downloaded at January 14, 2021.

**Table 1.** The number of PDB RNAs classified according to N*_eff_*-value obtained from RNAcmap alignment.

| $N_{eff}$-value | Total | Rfam mapped | Non-Rfam mapped |
| --- | --- | --- | --- |
| No-hit | 21 | 1 | 20 |
| Low ($1 \leq N_{eff} \leq 10$) | 83 | 15 | 68 |
| Medium ($10 < N_{eff} \leq 50$) | 31 | 15 | 16 |
| High ($50 < N_{eff}$) | 110 | 87 | 23 |
| Total | 245 | 118 | 127 |

### Rfam mapping and MSA

The above 245 PDB structures were mapped into Rfam and non-Rfam families by directly searching the PDB RNA sequence on the Rfam website (https://rfam.xfam.org/). Out of 245 chains, 118 RNA sequences were mapped to 48 different Rfam families, whereas 127 could not be mapped to any existing Rfam families, as shown in Table 1.

To obtain Rfam MSAs on Rfam mapped RNAs, we downloaded fasta files of 48 families from http://ftp.ebi.ac.uk/pub/databases/Rfam/14.6/fasta_files/ and corresponding covariance model files (Rfam.cm.gz) from http://ftp.ebi.ac.uk/pub/databases/Rfam/14.6/. The query sequence along with its corresponding family fasta sequences were aligned using *cmalign* program from INFERNAL along with the covariance model of that particular family as an input.

### Reference database

Both RNAcmap and RNAcmap2 require a reference database for homology search. In RNAcmap, we employed NCBI’s nucleotide (*nt*) database [17] as the only reference database library. In this work, we have expanded our reference database with environment samples (*env_nt*), transcriptome shotgun assembly (*tsa_nt*), and nucleotide sequences derived from the Patent Division of GenBank (*pat_nt*) databases in addition to NCBI’s nucleotide (nt) database.

The NCBI’s nucleotide database file (*nt.gz*) used in RNAcmap and RNAcmap2 was downloaded from https://ftp.ncbi.nlm.nih.gov/blast/db/FASTA/ on January 14, 2021. NCBI’s nucleotide database (nt) file was of size 344 GB after unzip. In addition to *nt* database, we downloaded *env_nt* (Version-1.1), *tsa_nt* (Version-1.1), and *pat_nt* (Version-1.1) databases from https://ftp.ncbi.nlm.nih.gov/blast/db/ on July 29, 2021. These four databases (*nt, env_nt, tsa_nt*, and *pat_nt*) were concatenated together using their fasta files into a single fasta file of size 506 GB. Duplicate sequences in the combined database were removed using the program SeqKit [23]. The final database is of size 487 GB used as a reference library for the RNAcmap2 pipeline. There is an increase 31% in the number of sequences for the expanded sequence library.

### The RNAcmap2 Pipeline

RNAcmap2 employs three iterative homology searches. There is one more INFERENAL search than RNAcmap, as shown in Figure 1. In the first search, a query RNA sequence is searched against the reference database library using BLAST-N [12] with *E-value=0.001, line-length=1000*, and *number-of-alignments=50000* to obtain multiple sequence alignment (MSA-1). Next, a covariance model (CM) is built from the MSA-1 and predicted secondary structure from RNAfold [1] [as consensus secondary structure (CSS)] using program *cmbuild* from the INFERNAL [15] tool. This CM is calibrated (Calibrated CM) using program *cmcalibrate* from the INFERNAL tool. Finally, the calibrated CM is searched against the reference database library using the *cmsearch* program from INFERNAL with *E-value=10* to obtain MSA-2.

**Fig. 1:**
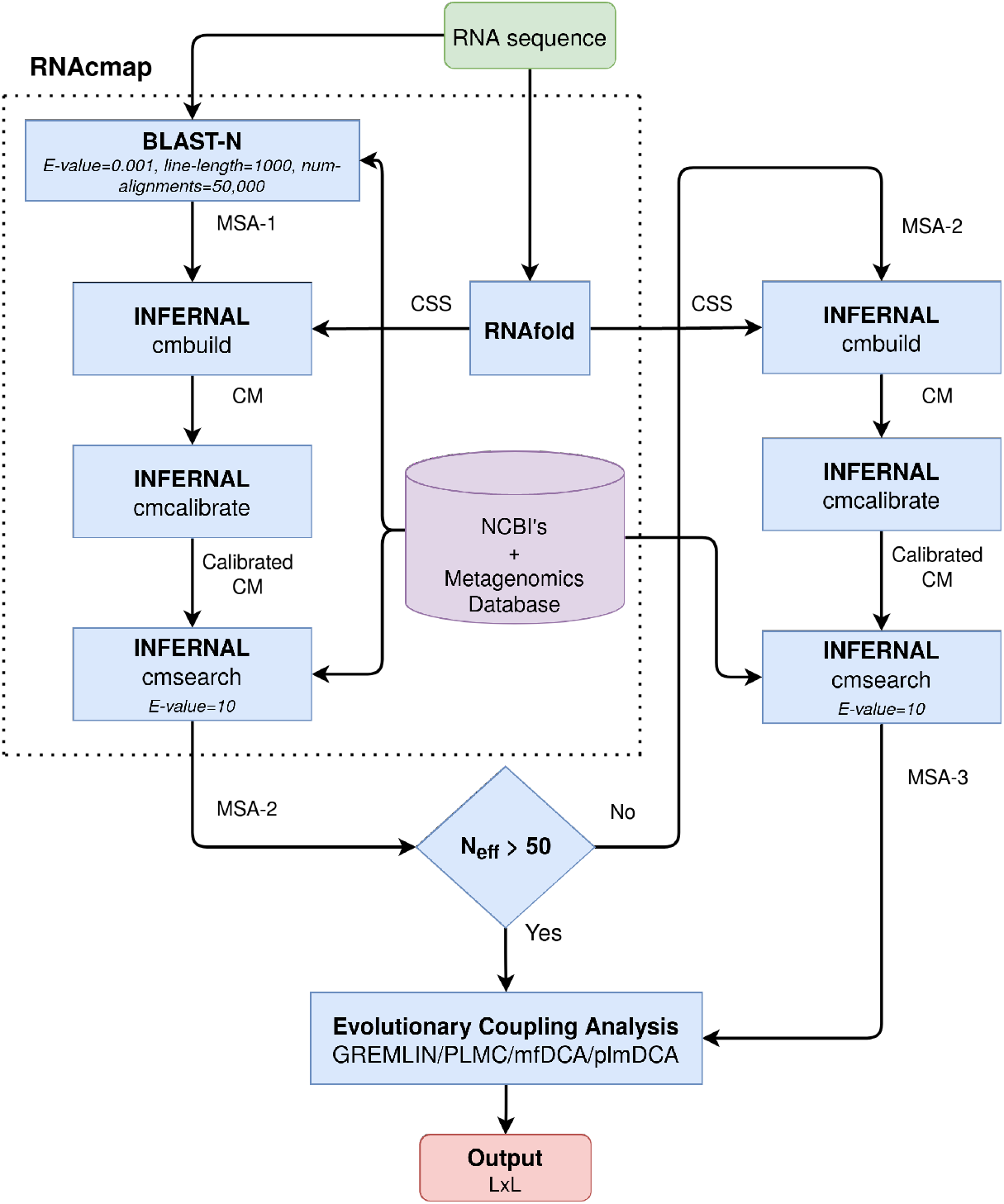
The architecture of the RNAcmap2 pipeline. CSS: Consensus Secondary Structure. CM: Covariance Model. L: Length of the input RNA sequence

If N*_eff_* <50 after the completion of the above two searches, we will perform the third round of homology search. This is done by building a new covariance model (CM) with the cmbuild program, which utilizes MSA-2 and the consensus secondary structure from RNAfold as an input. The new covariance model (CM) after calibration is then employed to search against the reference database library with the same e-value of 10 to obtain the MSA-3, as shown in Figure 1. MSA-3 was considered as the final set of homologous sequences. Here, we stop the search after two searches if N*_eff_* >50 because method performance does not improve much after N*_eff_* >50 in secondary structure and contact map prediction (SPOT-RNA2 [8] and SPOT-RNA-2D [24]). In this work, we found that three iterations were usually sufficient for most cases. However, additional iterations may be beneficial if extreme sensitivity is required and long computational time is not a concern.

### Performance Evaluation by Base Pairing

One way to evaluate MSAs generated from RNAcmap and RNAcmap2 is to compare the structural accuracy extracted from co-variational analysis of MSAs. This was done by employing direct coupling analysis predictors such as GREMLIN [3], PLMC [5, 6], mfDCA [7], and plmDCA [4]. GREMLIN and PLMC are pseudo-likelihood optimization-based DCA predictors. GREMLIN and PLMC were downloaded from https://github.com/sokrypton/GREMLIN_CPP and https://github.com/debbiemarkslab/plmc, respectively, and both run with the default parameters. Mean-field direct coupling analysis (mfDCA) and pseudo-likelihood maximization (plmDCA) direct coupling analysis algorithms were obtained from pydca [25] program (https://github.com/KIT-MBS/pydca).

Another way to evaluate the usefulness of homologous sequences obtained is to employ the alignment-based folding algorithm RNAalifold [1]. RNAalifold was obtained from Vienna package version 2.4.14 (https://www.tbi.univie.ac.at/RNA/). The program was run with the default parameters. As a comparison, we also obtained CaCofold [2], which uses both positive and negative evolutionary information along with a probabilistic folding algorithm for RNA secondary structure prediction. CaCofold was downloaded from http://eddylab.org/R-scape/ and was run with the default parameters.

The third way to evaluate homologous sequences is to input the resulting sequence profile and direct coupling result into our deep-learning-based predictor SPOT-RNA2 [8]. SPOT-RNA2 was downloaded from https://github.com/jaswindersingh2/SPOT-RNA2 and run with the default settings.

### Performance Evaluation by Structure Alignment

In addition to the performance evaluation according to secondary structure, one direct way to compare RNAcmap2 MSAs with other MSAs is by comparing the alignment accuracy of RNAs belonging to the same Rfam family and having 3D structures. We first located Rfam families that contain more than one RNA with known structure. Then the “gold-standard” alignment is made by performing structure alignment using RMalign [26]. To obtain the RNAcmap2 alignment, one RNA from each family with the highest N*_eff_* was considered as the reference RNA, and the covariance model was built by using RNAcmap2 MSA. Next, the remaining RNAs within a family were aligned with the highest N_*eff*_ RNA’s covariance model using the *cmscan* program from INFERNAL. To obtain the Rfam alignment, we used the covariance model provided by the Rfam for that particular family to align RNAs within the same family. The alignment accuracy is measured by comparing Rfam and RNAcmap2 generated alignments to the gold standard structure alignment.

### Performance Measures

For benchmarking four DCA and alignment-based folding predictors on RNAcmap’s MSAs, sensitivity (*SN* = *TP*/(*TP* + *FN*)), precision (*PR* = *TP*/(*TP* + *FP*)), and F1-score [F1 = 2(*PR* × *SN*)/(*PR* + *SN*)] were used for non-local base-pairs [|*i* – *j*| ≥ 4], where *TP*, *TN*, *FP* and *FN* are true positives, true negatives, false positives and false negatives, respectively. The performance metrics were evaluated for individual RNAs with mean performance reported in the results section.

## Results

### Improvement of MSAs measured by coupling analysis

The quality of MSAs generated by different methods can be quantified by the accuracy of base pairs inferred from direct coupling analysis. Here, we employed the mfDCA predictor as other predictors yield the same trend, and mfDCA has the best performance, as shown in Supplementary Table S1. Table 2 (the average result) and Figure 2 (the distribution) compare three MSAs generated by BLAST-N, direct INFERNAL, the combination of BLAST-N and INFERNAL (RNAcmap) searches against the NCBI nt database along with two additional MSAs by the RNAcmap (RNAcmap*) and RNAcmap2 searches against the expanded NCBI database (nt, env nt, tsa nt, and pat nt). The performance measure is the F1-score for top L/3 predictions.

**Table 2.**
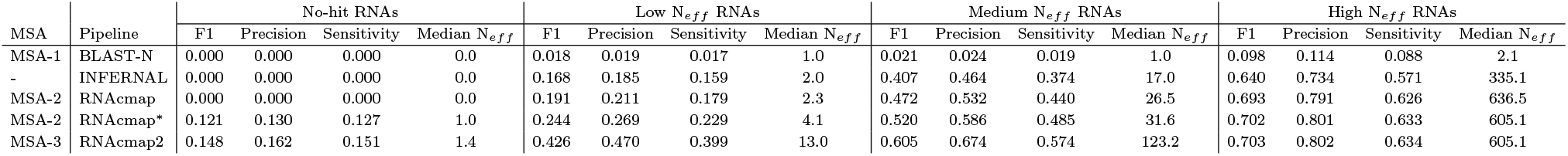
Performance comparison among different MSA pipelines on No-hit RNAs (21 RNAs), Low N*_eff_* RNAs (83 RNAs), Medium N*_eff_* RNAs (31 RNAs), and High N*_eff_* RNAs (110 RNAs) using mfDCA predictor.

**Fig. 2:**
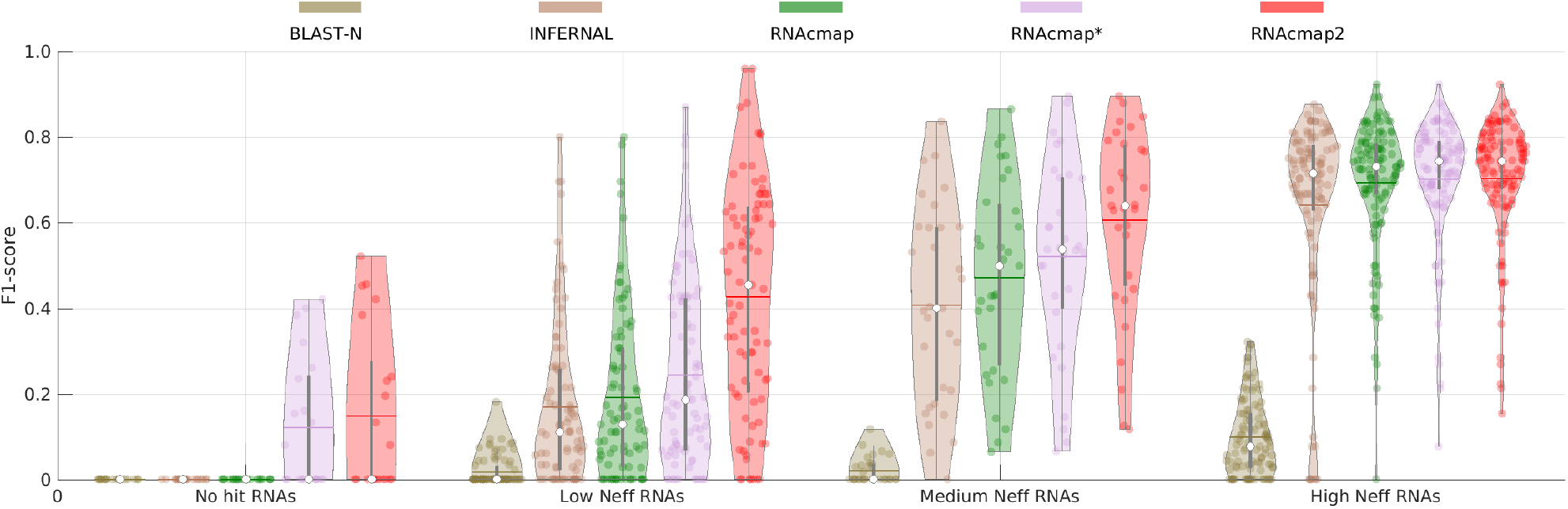
Violin plot of F1-score of predicted top L/3 contacts by mfDCA predictor using BLAST-N, INFERNAL, RNAcmap, RNAcmap*, and RNAcmap2 supplied aligned homologous sequences for No-hit RNAs (21 RNAs), Low *N_eff_* RNAs (83 RNAs), Medium *N_eff_* RNAs (31 RNAs) and high *N_eff_* RNAs (110 RNAs). In the Violin plot: white dot represents the median; the thin horizontal line represents mean; the thick and thin gray bar in the center represents the interquartile range and 1.5×interquartile range; the curve on either side of gray line shows the distribution of data using kernel density estimation; wider the curves around gray lines, higher the probability of data points lies in that region and vice versa.

As Table 2 and Figure 2 show, BLAST-N is quite ineffective in getting a significant number of effective homologs for all RNAs (F1-scores all <0.1), confirming that sequences are not very conserved in RNA homologs. Direct INFERNAL search with secondary structure predicted by RNAfold allows substantial improvement over BLAST-N for low, medium and N*_eff_* RNAs. Combining BLAST-N and INFERNAL (RNAcmap) provides an additional 14%, 16%, and 10% improvement over INFERNAL for these low, medium and high-N*_eff_* RNAs, respectively. Expansion of the sequence library leads to a 10% increase in F1-score for medium N*_eff_* RNAs and a 27% increase for low N_*eff*_ RNAs from RNAcmap to RNAcmap*. An additional round of homology search leads to a 16% increase in F1-score for medium *N_eff_* RNAs and a 75% increase for low *N_eff_* RNAs from RNAcmap* to RNAcmap2. For no-hit RNAs, the expansion of the sequence library allows detection of some homologs (Median *N_eff_*= 1) by RNAcmap* with further improvement by RNAcmap2 (Median *N_eff_*= 1.4). Similar trends were obtained from the other three DCA predictors as shown in Supplementary Tables S2, S3, and S4.

One interesting observation is that median *N_eff_*-values for high *N_eff_* RNAs obtained from RNAcmap (*N_eff_*=636.5) was slightly higher than RNAcmap2 (*N_eff_*=605.1). This is counter-intuitive as RNAcmap employed a reference database that is a subset of the RNAcmap2 reference database. This is due to the limit of top 50,000 allowed hits. If top 50,000 contains more redundant sequences, the number of effective homologous sequences will be lower. Indeed, if we relax 50,000 to 100,000, we will achieve higher *N_eff_* for RNAcmaps. However, we keep 50,000 as the default to save the computational time.

To illustrate the consistent improvement of RNAcmap2 over RNAcmap, Figure 3 compares the performance of RNAcmap2 and RNAcmap supplied alignments at different cutoffs for top predictions. RNAcmap2 improves over RNAcmap at all cutoffs for top contact predictions (L/n, n=1,2,3,…9,10). The results are similar for all other three DCA predictors (See Supplementary Figures S1, S2, and S3).

**Fig. 3:**
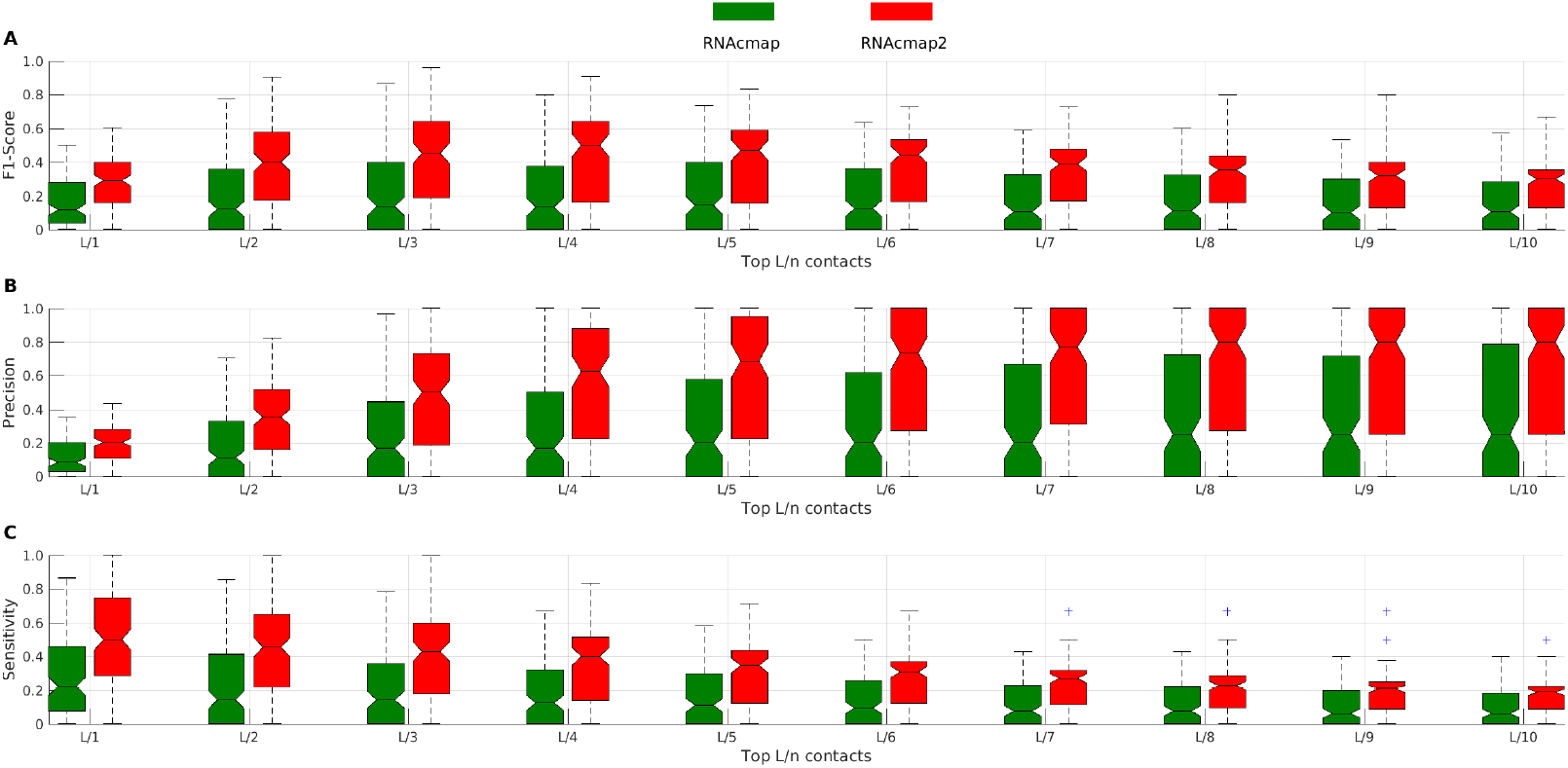
Boxplot of F1-score (**A**), Precision (**B**), and Sensitivity (**C**) as a function of predicted top L/n base pairs by mfDCA from RNAcmap (in green) and RNAcmap2 (in red) supplied alignment for 135 PDB RNAs from no-hit, low, medium N*_eff_* test sets. The distribution is shown in terms of median, 25th and 75th percentile with outlier shown by dots.

### RNAcmap2 vs Rfam

Rfam is considered the de facto gold standard for clustering homologous RNA families and their multiple sequence alignments because they have employed experimental secondary structure for alignment and homology search where possible. From 245 RNAs, 118 RNAs were mapped to Rfam families, while the remaining 127 RNAs were not mapped to any Rfam family (See Table 1). Figure 4 compares F1-scores of top L/3 predicted base pairs by mfDCA on MSAs from Rfam and that from RNAcmap2 for those RNAs mapped to Rfam [No-hit (1 RNA) and Low N*_eff_* (15 RNAs), Medium N*_eff_* (15 RNAs), and High N*_eff_* (87 RNAs) test sets]. The fully automatic RNAcmap2 MSAs yield essentially the same performance on base-pair prediction as the manually curated Rfam for high N*_eff_* RNAs and improve it over somewhat for low N*_eff_* RNAs and significantly for medium N*_eff_* RNAs.

**Fig. 4:**
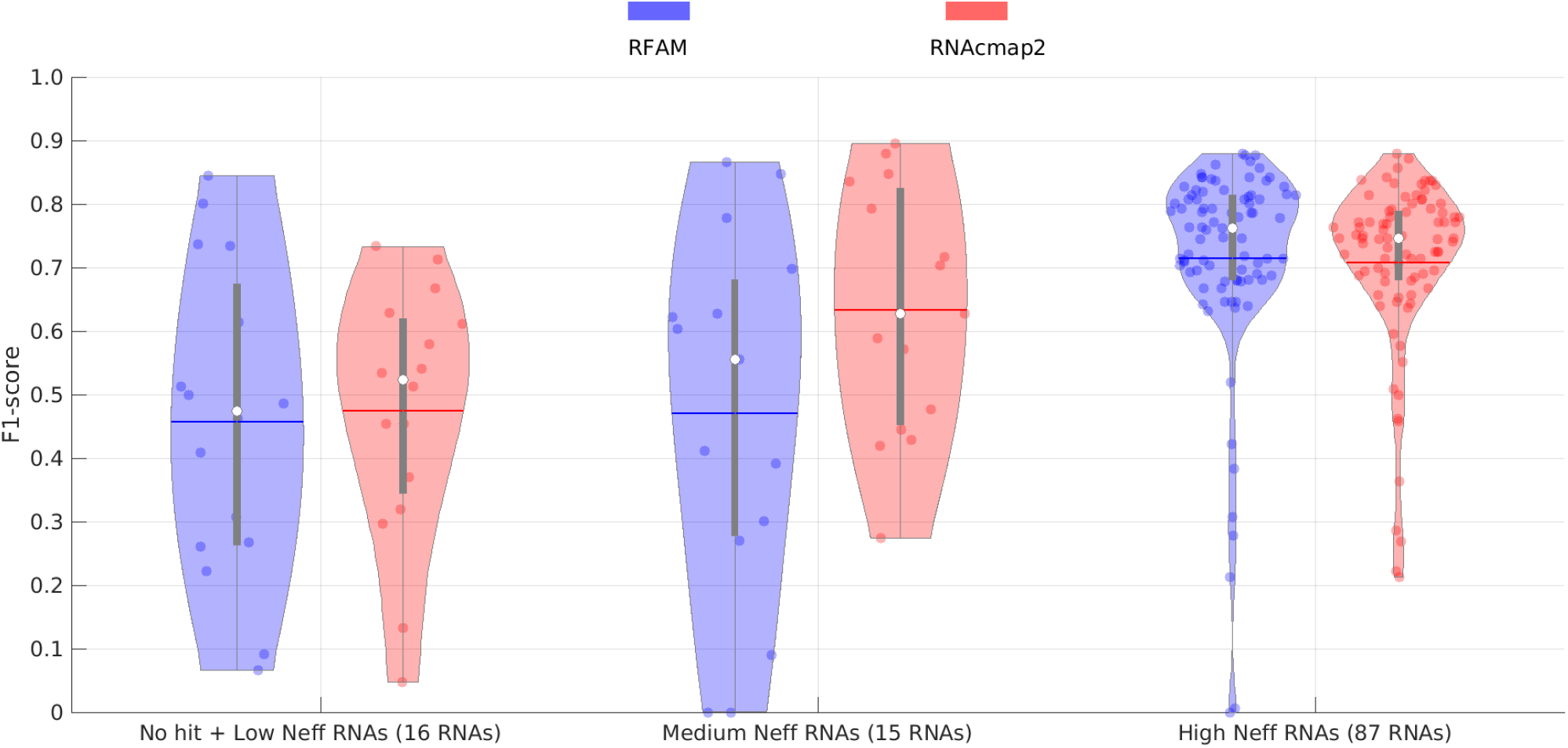
F1-score of predicted top L/3 base pairs by mfDCA from Rfam (in blue) and RNAcmap2 (in red) supplied alignment for No-hit RNAs (1 RNA), Low N*_eff_* RNAs (15 RNAs), Medium N*_eff_* RNAs (15 RNAs) and high N*_eff_* RNAs (87 RNAs) mapped to Rfam. In the Violin plot: white dot represents the median; the thin horizontal line represents mean; the thick and thin gray bar in the center represents the interquartile range and 1.5×interquartile range; the curve on either side of gray line shows the distribution of data using kernel density estimation; wider the curves around gray lines, higher the probability of data points lies in that region and vice versa.

Table 3 shows a family-wise performance comparison of RNAcmap2, Rfam, and RNAcmap supplied MSAs using top L/3 mfDCA predicted base pairs on 48 different Rfam families. RNAcmap2 MSAs performed better than Rfam MSAs on 29/48 families, and Rfam MSAs performed better than RNAcmap2 for 16/48 families, while performance on three families was equally good.

**Table 3.** Performance comparison of RNAcmap, Rfam, and RNAcmap2 based on F1-score on 48 Rfam mapped families using mfDCA top L/3 predicted base pairs.

| Rfam Family | RNA type | No. of RNAs | RNAmcap | Rfam | RNAmcap2 |
| --- | --- | --- | --- | --- | --- |
| RF00005 | tRNA | 53 | 0.717 | **0.746** | 0.724 |
| RF00001 | 5S ribosomal | 6 | 0.618 | **0.630** | 0.625 |
| RF00167 | Purine riboswitch | 4 | **0.711** | 0.685 | **0.711** |
| RF01852 | Selenocysteine transfer | 3 | 0.856 | 0.852 | **0.871** |
| RF00379 | ydaO/yuaA leader | 3 | 0.568 | **0.693** | 0.597 |
| RF01750 | ZMP/ZTP riboswitch | 2 | 0.406 | **0.657** | 0.511 |
| RF00162 | SAM riboswitch | 2 | **0.757** | 0.746 | **0.757** |
| RF00234 | glmS glucosamine-6-phosphate activated ribozyme | 2 | **0.709** | 0.695 | 0.697 |
| RF01051 | Cyclic di-GMP-I riboswitch | 2 | 0.647 | **0.704** | 0.647 |
| RF00059 | TPP riboswitch | 2 | 0.453 | **0.679** | 0.485 |
| RF00174 | Cobalamin riboswitch | 2 | **0.650** | 0.528 | 0.640 |
| RF01689 | AdoCbl variant | 1 | 0.000 | **0.613** | 0.453 |
| RF02680 | PreQ1-III riboswitch | 1 | 0.029 | 0.261 | **0.319** |
| RF00390 | UPSK | 1 | 0.080 | 0.066 | **0.297** |
| RF02679 | Pistol ribozyme | 1 | 0.111 | 0.486 | **0.541** |
| RF02519 | ToxI antitoxin | 1 | 0.364 | 0.091 | **0.455** |
| RF01734 | Fluoride riboswitch | 1 | 0.171 | **0.800** | 0.629 |
| RF02796 | Pab160 | 1 | 0.462 | 0.462 | **0.667** |
| RF01763 | Guanidine-III riboswitch | 1 | 0.462 | **0.513** | **0.513** |
| RF01704 | Downstream peptide | 1 | 0.667 | **0.733** | **0.733** |
| RF03013 | nadA | 1 | 0.263 | **0.737** | 0.579 |
| RF01725 | SAM-I/IV variant riboswitch | 1 | 0.000 | 0.500 | **0.611** |
| RF01415 | Flavivirus 3'UTR stem loop IV | 1 | 0.222 | **0.267** | 0.133 |
| RF00102 | VA | 1 | 0.073 | 0.409 | **0.713** |
| RF00100 | 7SK | 1 | 0.400 | **0.844** | 0.533 |
| RF02678 | Hatchet ribozyme | 1 | 0.259 | 0.222 | **0.370** |
| RF00026 | U6 spliceosomal | 1 | 0.143 | 0.000 | **0.429** |
| RF01826 | SAM-V riboswitch | 1 | 0.244 | 0.300 | **0.476** |
| RF00442 | Guanidine-I riboswitch | 1 | 0.232 | **0.556** | 0.274 |
| RF02553 | Y RNA-like | 1 | 0.254 | **0.698** | 0.419 |
| RF00505 | RydC | 1 | 0.432 | 0.270 | **0.571** |
| RF01344 | CRISPR RNA direct repeat element | 1 | 0.429 | 0.000 | **0.880** |
| RF02683 | NiCo riboswitch | 1 | 0.394 | 0.627 | **0.836** |
| RF00458 | Cripavirus internal ribosome entry site | 1 | 0.500 | 0.391 | **0.628** |
| RF00164 | Coronavirus 3' stem-loop II-like motif | 1 | **0.647** | 0.412 | 0.588 |
| RF00008 | Hammerhead ribozyme | 1 | 0.755 | 0.604 | **0.792** |
| RF01786 | Cyclic di-GMP-II riboswitch | 1 | 0.542 | **0.847** | **0.847** |
| RF00228 | Hepatitis A virus internal ribosome entry site | 1 | 0.418 | 0.090 | **0.716** |
| RF01831 | THF riboswitch | 1 | 0.600 | **0.714** | 0.657 |
| RF01854 | Bacterial large signal recognition particle | 1 | 0.595 | **0.762** | 0.643 |
| RF02001 | Group II catalytic intron D1-D4-3 | 1 | **0.726** | 0.006 | 0.720 |
| RF00050 | FMN riboswitch | 1 | 0.698 | 0.721 | **0.744** |
| RF01857 | Archaeal signal recognition particle | 1 | 0.793 | 0.793 | **0.811** |
| RF00029 | Group II catalytic intron | 1 | **0.760** | 0.520 | **0.760** |
| RF00168 | Lysine riboswitch | 1 | **0.752** | 0.709 | **0.752** |
| RF00504 | Glycine riboswitch | 1 | **0.831** | 0.800 | **0.831** |
| RF00380 | M-box riboswitch | 1 | **0.677** | 0.632 | 0.647 |
| RF00002 | 5.8S ribosomal | 1 | 0.378 | **0.422** | 0.222 |
| Mean | 48 Families | - | 0.468 | 0.531 | **0.605** |
| Non-Rfam | - | 127 | 0.278 | - | **0.440** |

Another way to compare Rfam MSAs with RNAcmap2 MSAs is to compare alignment accuracy. Table 4 shows that the overall performance is nearly the same. RNAcmap2 has a better alignment on 6/11 families whereas Rfam has a better alignment on 5/11 families with two or more RNA structures. The median accuracies are 78.89% for RNAcmap2 and 78.38% for Rfam, respectively.

**Table 4.** Comparison of alignment accuracy given by Rfam and RNAcmap2 within a family with respect to structure alignment obtained from RMalign using tertiary structures.

| Rfam Family | RNA type | No. of RNAs | Mean Accuracy |  |
| --- | --- | --- | --- | --- |
|  |  |  | Rfam | RNAmcp2 |
| RF00001 | 5S ribosomal | 5 | 76.20 | **89.56** |
| RF00005 | tRNA | 52 | **82.27** | 71.93 |
| RF00059 | TPP riboswitch | 1 | **90.79** | 89.47 |
| RF00162 | SAM riboswitch | 1 | 82.98 | **89.36** |
| RF00167 | Purine riboswitch | 3 | **94.81** | 94.29 |
| RF00174 | Cobalamin riboswitch | 1 | **69.81** | 45.28 |
| RF00234 | glmS glucosamine-6-phosphate activated ribozyme | 1 | 68.64 | **73.73** |
| RF00379 | ydaO/yuaA leader | 2 | 78.38 | **78.89** |
| RF01051 | Cyclic di-GMP-I riboswitch | 1 | 88.00 | **93.33** |
| RF01750 | ZMP/ZTP riboswitch | 1 | **67.39** | 65.22 |
| RF01852 | Selenocysteine transfer | 2 | 59.18 | **60.94** |
| Mean | - | - | **78.04** | 77.45 |
| Median | - | - | 78.38 | **78.89** |
| Median $N_{eff}$ | - | - | **3206.19** | 710.48 |

### Application of RNAcmap2 MSA

RNAcmap2 MSAs can be employed for improving different applications such as DCA predictors (mfDCA), alignment-based folding predictors (RNAaliFold and CaCoFold) and deep-learning-based predictors (SPOT-RNA2). Figure 5**A** shows the method performance for a combined 38 RNAs from no-hit, low, median *N_eff_* sets and 37 RNAs from high *N_eff_* set with ‘deep’ RNAcmap2 alignment (*N_eff_*/L > 0.2) after excluding sequences overlapping with SPOT-RNA2 training data using CD-HIT-EST at lowest identity cut-off of 0.8.

**Fig. 5:**
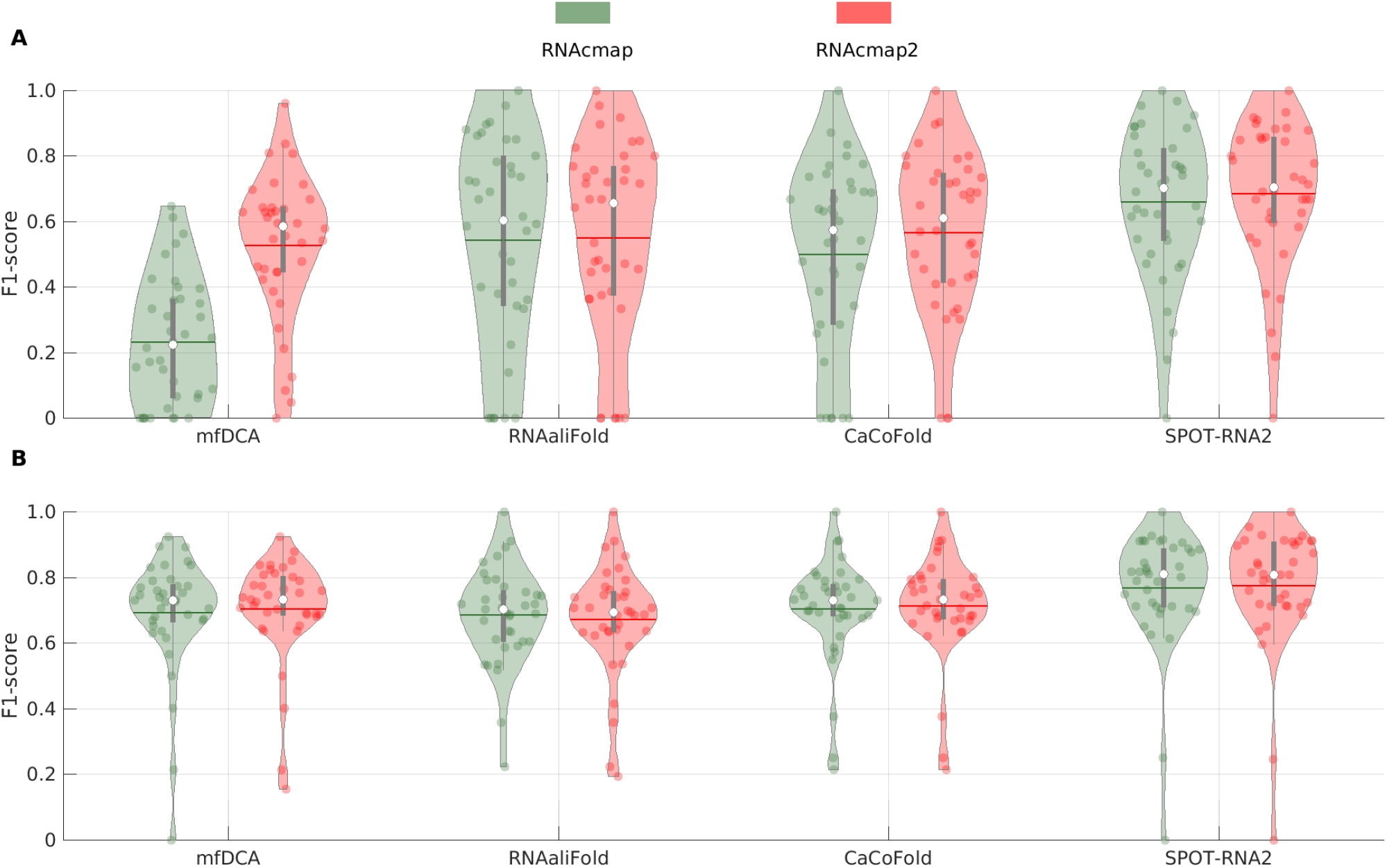
Violin plot of F1-score of base pairs (bps) predictions from mfDCA, RNAaliFold, CaCoFold, and SPOT-RNA2 using RNAcmap and RNAcmap2 supplied alignment for combined no-hit, low, and median N*_eff_* RNAs (**A**) and high N*_eff_* RNAs (**B**). The metrics are evaluated on combined 38 RNAs from no-hit, low, median N*_eff_* sets and 37 RNAs from high N*_eff_* set with ‘deep’ RNAcmap2 alignment (N*_eff_*/L > 0.2) and non-redundant from SPOT-RNA2 training data at 80% identity cut-off according to CD-HIT-EST. In the Violin plot: white dot represents the median; the thin horizontal line represents mean; the thick and thin gray bar in the center represents the interquartile range and 1.5×interquartile range; the curve on either side of gray line shows the distribution of data using kernel density estimation; wider the curves around gray lines, higher the probability of data points lies in that region and vice versa.

RNAcmap2 MSA provides significant improvement for base pairs prediction over RNAcmap MSA in all cases with the largest improvement for mfDCA as it has the lowest performance with RNAcmap MSA. The deep learning SPOT-RNA2 has the smallest improvement because it is more difficult to improve over already accurate prediction with the median F1-score >0.70 for SPOT-RNA2 with the RNAcmap input, while all other predictors with F1-score <0.66 even after using RNAcmap2 MSAs.

For high *N_eff_* RNAs (Figure 5**B**), RNAcmap2 supplied MSAs performance are essentially the same as the RNAcmap MSAs, confirming the use of N*_eff_*=50 as the cutoff for stopping the search.

As an illustrative example, Figure 6 compares RNAcmap and RNAcmap2 MSAs based on base pair prediction by mfDCA and SPOT-RNA2. This is tRNA in a protein-tRNA complex structure (chain B in PDB ID 4wj4) [27]. In this figure, correctly predicted canonical base-pairs, non-canonical base-pairs and pseudoknots are indicated by the color blue, orange and green, respectively. Any wrongly predicted base-pairs are shown by color magenta. mfDCA (RNAcmap2) predicted a more accurate native-like topology (Figure 6A) with an F1-score of 0.81 as compared to mfDCA (RNAcmap) with an F1-score of 0.77 (Figure 6B) in comparison to the native structure (Figure 6C). In both cases, mfDCA is able to predict one pseudoknot (in green) and non-canonical (in orange) base-pair correctly. SPOT-RNA2 (RNAcmap2) is able to predict all the canonical, non-canonical, and pseudoknot base-pairs in the native structure with an overall native-like topology as shown in Figure 6E and F1-score of 0.93. In comparison, SPOT-RNA2 (RNAcmap) predicts structure with an F1-score of 0.82 as shown in Figure 6D, but due to many predicted false base-pairs, the overall topology of the predicted structure by SPOT-RNA2 (RNAcmap) is quite different from the native structure in Figure 6F. The result highlights the power of improved MSAs for secondary structure and tertiary base-pair prediction.

**Fig. 6:**
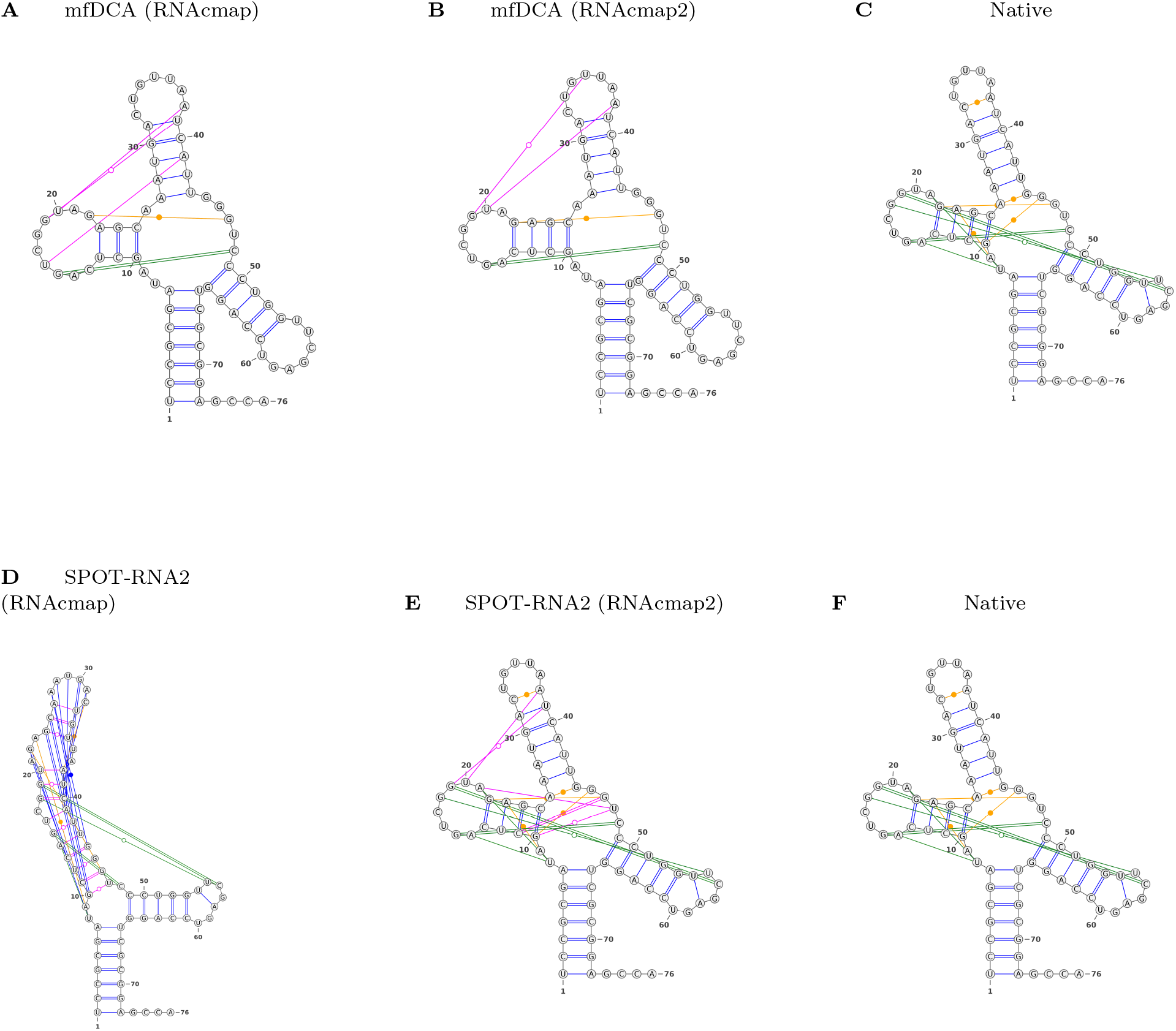
Performance illustrated by a tRNA (chain B in PDB ID 4wj4). **(A)** Predicted secondary structure by mfDCA using RNAcmap alignment, with F1-score of 77%. **(B)** Predicted secondary structure by mfDCA using RNAcmap2 alignment, with F1-score of 81%. **(C)** Native structure. **(D)** Predicted secondary structure by SPOT-RNA2 using RNAcmap alignment, with F1-score of 82%. **(E)** Predicted secondary structure by SPOT-RNA2 using RNAcmap2 alignment, with F1-score of 93%. **(F)** Native structure.

## Discussion

This work has developed a new automatic homology search pipeline (RNAcmap2) by performing three iterative searches against an expanded sequence library, including metagenomics and patented sequences. RNAcmap2 significantly improved over RNAcmap with mean F1-score increased from 0.00 to 0.148, 0.191 to 0.426, and 0.472 to 0.605 for No-hit, low N*_eff_*, and medium N*_eff_* RNAs, respectively. Moreover, RNAcmap2 produced MSAs that are comparably accurate as manually curated Rfam MSAs when compared on 118 PDB RNAs belonging to 48 Rfam families in their ability to extract base pairing information. Thus, RNAcmap2 can generate MSAs for any Rfam or non-Rfam RNAs with the Rfam-like performance. The new tool is expected to be useful for predicting 1D-structural properties such as solvent accessibility [9, 10, 11] and backbone torsion angle prediction [28], 2D structural properties such as base pairing [8] and contact maps [24, 29], and RNA 3D structure prediction [30, 31, 32].

The performance of RNAcmap2 depends on the accuracy of Consensus Secondary Structure (CSS). Here, we employed RNAfold as it is one of the state-of-the-art, folding-based algorithms for RNA secondary structure prediction. Recently, deep learning techniques such as single-sequence-based SPOT-RNA significantly improves over folding-based algorithms in predicting RNA secondary structures [33]. Supplementary Table S5 shows the mfDCA performance comparison of RNAcmap* and RNAcmap2 with RNAfold and SPOT-RNA as the CSS predictor on 102 RNAs after removing redundant sequences to SPOT-RNA training set using CD-HIT-EST at 80% identity cut-off. RNAcmap* with SPOT-RNA CSS leads to 9% higher F1-score over RNAcmap* with RNAfold CSS.Similarly, RNAcmap2 with SPOT-RNA CSS yields 3% higher F1-score than the RNAcmap2 (RNAfold). In the automatic server provided, we give users an option for using RNAfold or SPOT-RNA for CSS. We did not choose SPOT-RNA as the default to avoid the potential over-fit issue.

RNAcmap2 is computationally expensive. For low N*_eff_* RNAs, RNAcmap2 takes about 5 hours for a 200 nts sequence when allowed to use 16 CPU threads of Intel(R) Xeon(R) CPU E5-2670 (2.60 GHz). Searching homologs for RNAs with >1000 nts is computationally prohibitive. This is why we restricted the RNAcmap2 pipeline for three iterative searches only as the default. However, if one has the necessary computational resource, the additional number of iterations can generate more homologous sequences. For no-hit RNAs, we will get a significant 9% improvement for the F1-score (0.148 to 0.162) from the mfDCA predictor for the 4^*th*^ round of the homology search. In the standalone version of RNAcmap2, the default value for the number of iterations can be set by users.

Despite the large improvement from RNAcmap to RNAcmap2, there are still 8 RNAs with PDB structures having no hits. RNAcmap2 can improve over RNAcmap only if there is more than one hit in MSA-2. This limitation will be overcome by the exponential increase in collection of RNA sequences [17].

## Key Points

- We developed an improved version of the fully automated RNA homology search pipeline (RNAcmap) by utilizing one BLAST-N search and two additional INFERNAL searches.
- RNAcmap2 expanded the reference NCBI’s nucleotide database by including NCBI’s metagenomics and patent sequences and improved the performance over RNAcmap for low N*_efff_* RNAs.
- We provide a publicly available standalone program at GITHUB and web-server on SPARKS-LAB.

## Availability

The RNAcmap2 pipeline is available at https://sparks-lab.org/server/rnacmap2/. It is highly recommended that users install the standalone program at https://github.com/jaswindersingh2/RNAcmap2.

## Author contributions statement

JS (Jaswinder Singh) and KP designed pipeline architecture. JS (Jaswinder Singh) and JS prepared the data. JS (Jaswinder Singh) and TL build reference database. JS (Jaswinder Singh) did the pipeline analysis, compared it with other pipelines and wrote the manuscript. JS (Jaswinder Singh) build the standalone program and webserver of the pipeline. YZ conceived the study, participated in the initial design, assisted in data analysis, and drafted the whole manuscript. All authors read, contributed to the discussion, and approved the final manuscript.

## Acknowledgments

We gratefully acknowledge the use of the High-Performance Computing Cluster Gowonda to complete this research and the aid of the research cloud resources provided by the Queensland Cyber Infrastructure Foundation (QCIF). We also gratefully acknowledge the support of NVIDIA Corporation with the donation of the Titan V GPU used for this research. This work is also supported in part by the High Performance Computing Cluster at Shenzhen Bay Laboratory. The support of Shenzhen Science and Technology Program (Grant No.KQTD20170330155106581) and the Major Program of Shenzhen Bay Laboratory S201101001 is also acknowledged.

## References

1. Ronny Lorenz, Stephan H. Bernhart, Christian Höner zu Siederdissen, Hakim Tafer, Christoph Flamm, Peter F. Stadler, and Ivo L. Hofacker. ViennaRNA Package 2.0. Algorithms for Molecular Biology, 6(1):26, 2011.

2. Elena Rivas. RNA structure prediction using positive and negative evolutionary information. PLOS Computational Biology, 16(10):1–25, 2020.

3. Hetunandan Kamisetty, Sergey Ovchinnikov, and David Baker. Assessing the utility of coevolution-based residue–residue contact predictions in a sequence- and structure-rich era. Proceedings of the National Academy of Sciences, 110(39):15674–15679, 2013.

4. Magnus Ekeberg, Cecilia Lövkvist, Yueheng Lan, Martin Weigt, and Erik Aurell. Improved contact prediction in proteins: Using pseudolikelihoods to infer Potts models. Phys. Rev. E, 87:012707, 2013.

5. Sivaraman Balakrishnan, Hetunandan Kamisetty, Jaime G. Carbonell, Su-In Lee, and Christopher James Langmead. Learning generative models for protein fold families. Proteins: Structure, Function, and Bioinformatics, 79(4):1061–1078, 2011.

6. Thomas A. Hopf, John B. Ingraham, Frank J. Poelwijk, Charlotta P. I. Schärfe, Michael Springer, Chris Sander, and Debora S. Marks. Mutation effects predicted from sequence co-variation. Nature Biotechnology, 35(2):128–135, 2017.

7. Faruck Morcos, Andrea Pagnani, Bryan Lunt, Arianna Bertolino, Debora S. Marks, Chris Sander, Riccardo Zecchina, José N. Onuchic, Terence Hwa, and Martin Weigt. Direct-coupling analysis of residue coevolution captures native contacts across many protein families. Proceedings of the National Academy of Sciences, 108(49):E1293–E1301, 2011.

8. Jaswinder Singh, Kuldip Paliwal, Tongchuan Zhang, Jaspreet Singh, Thomas Litfin, and Yaoqi Zhou. Improved RNA Secondary Structure and Tertiary Base-pairing Prediction using Evolutionary Profile, Mutational Coupling and Two-dimensional Transfer Learning. Bioinformatics, 37(17):2589–2600, 2021.

9. Yuedong Yang, Xiaomei Li, Huiying Zhao, Jian Zhan, Jihua Wang, and Yaoqi Zhou. Genome-scale characterization of RNA tertiary structures and their functional impact by RNA solvent accessibility prediction. RNA, 23(1):14–22, 2017.

10. Saisai Sun, Qi Wu, Zhenling Peng, and Jianyi Yang. Enhanced prediction of RNA solvent accessibility with long short-term memory neural networks and improved sequence profiles. Bioinformatics, 35(10):1686–1691, 2018.

11. Anil Kumar Hanumanthappa, Jaswinder Singh, Kuldip Paliwal, Jaspreet Singh, and Yaoqi Zhou. Single-sequence and profile-based prediction of RNA solvent accessibility using dilated convolutional neural network. Bioinformatics, 36(21):5169–5176, 2020.

12. Stephen F. Altschul, Thomas L. Madden, Alejandro A. Schäffer, Jinghui Zhang, Zheng Zhang, Webb Miller, and David J. Lipman. Gapped BLAST and PSI-BLAST: a new generation of protein database search programs. Nucleic Acids Research, 25(17):3389–3402, 1997.

13. T.F. Smith and M.S. Waterman. Identification of common molecular subsequences. Journal of Molecular Biology, 147(1):195–197, 1981.

14. Travis J. Wheeler and Sean R. Eddy.nhmmer: DNA homology search with profile HMMs. Bioinformatics, 29(19):2487–2489, 2013.

15. Eric P. Nawrocki and Sean R. Eddy. Infernal 1.1: 100-fold faster RNA homology searches. Bioinformatics, 29(22):2933–2935, 2013.

16. Ioanna Kalvari, Joanna Argasinska, Natalia Quinones-Olvera, Eric P Nawrocki, Elena Rivas, Sean R Eddy, Alex Bateman, Robert D Finn, and Anton I Petrov. Rfam 13.0: shifting to a genome-centric resource for non-coding RNA families. Nucleic Acids Research, 46(D1):D335–D342, 2017.

17. NCBI Resource Coordinators. Database resources of the National Center for Biotechnology Information. Nucleic Acids Research, 46(D1):D8–D13, 2017.

18. Tongchuan Zhang, Jaswinder Singh, Thomas Litfin, Jian Zhan, Kuldip Paliwal, and Yaoqi Zhou. RNAcmap: A Fully Automatic Pipeline for Predicting Contact Maps of RNAs by Evolutionary Coupling Analysis. Bioinformatics, 37(20):3494–3500, 2021.

19. Peter W. Rose, Andreas Prlić, Ali Altunkaya, Chunxiao Bi, Anthony R. Bradley, Cole H. Christie, Luigi Di Costanzo, Jose M. Duarte, Shuchismita Dutta, Zukang Feng, Rachel Kramer Green, David S. Goodsell, Brian Hudson, Tara Kalro, Robert Lowe, Ezra Peisach, Christopher Randle, Alexander S. Rose, Chenghua Shao, Yi-Ping Tao, Yana Valasatava, Maria Voigt, John D. Westbrook, Jesse Woo, Huangwang Yang, Jasmine Y. Young, Christine Zardecki, Helen M. Berman, and Stephen K. Burley. The RCSB protein data bank: integrative view of protein, gene and 3D structural information. Nucleic Acids Research, 45(D1):D271–D281, 2016.

20. Peter J. A. Cock, Tiago Antao, Jeffrey T. Chang, Brad A. Chapman, Cymon J. Cox, Andrew Dalke, Iddo Friedberg, Thomas Hamelryck, Frank Kauff, Bartek Wilczynski, and Michiel J. L. de Hoon. Biopython: freely available Python tools for computational molecular biology and bioinformatics. Bioinformatics, 25(11):1422–1423, 2009.

21. Limin Fu, Beifang Niu, Zhengwei Zhu, Sitao Wu, and Weizhong Li. CD-HIT: accelerated for clustering the next-generation sequencing data. Bioinformatics, 28(23):3150–3152, 2012.

22. Xiang-Jun Lu, Harmen J. Bussemaker, and Wilma K. Olson. DSSR: an integrated software tool for dissecting the spatial structure of RNA. Nucleic Acids Research, 43(21):e142–e142, 2015.

23. Wei Shen, Shuai Le, Yan Li, and Fuquan Hu. SeqKit: A Cross-Platform and Ultrafast Toolkit for FASTA/Q File Manipulation. PLOS ONE, 11(10):1–10, 2016.

24. Jaswinder Singh, Kuldip Paliwal, Thomas Litfin, Jaspreet Singh, and Yaoqi Zhou. Predicting RNA distance-based contact maps by integrated deep learning on physics-inferred secondary structure and evolutionary-derived mutational coupling. Bioinformatics, (Under-Review), 2022.

25. Mehari B Zerihun, Fabrizio Pucci, Emanuel K Peter, and Alexander Schug. pydca v1.0: a comprehensive software for direct coupling analysis of RNA and protein sequences. Bioinformatics, 36(7):2264–2265, 2019.

26. Jinfang Zheng, Juan Xie, Xu Hong, and Shiyong Liu. RMalign: an RNA structural alignment tool based on a novel scoring function RMscore. BMC Genomics, 20(1):276, 2019.

27. Tateki Suzuki, Akiyoshi Nakamura, Koji Kato, Dieter Söll, Isao Tanaka, Kelly Sheppard, and Min Yao. Structure of the pseudomonas aeruginosa transamidosome reveals unique aspects of bacterial trna-dependent asparagine biosynthesis. Proceedings of the National Academy of Sciences, 112(2):382–387, 2015.

28. Jaswinder Singh, Kuldip Paliwal, Jaspreet Singh, and Yaoqi Zhou. RNA Backbone Torsion and Pseudotorsion Angle Prediction Using Dilated Convolutional Neural Networks. Journal of Chemical Information and Modeling, 61(6):2610–2622, 2021.

29. Saisai Sun, Wenkai Wang, Zhenling Peng, and Jianyi Yang. RNA inter-nucleotide 3D closeness prediction by deep residual neural networks. Bioinformatics, 37(8):1093–1098, 2020.

30. Eleonora De Leonardis, Benjamin Lutz, Sebastian Ratz, Simona Cocco, Rémi Monasson, Alexander Schug, and Martin Weigt. Direct-Coupling Analysis of nucleotide coevolution facilitates RNA secondary and tertiary structure prediction. Nucleic acids research, 43(21):10444–10455, 2015.

31. Jian Wang, Kangkun Mao, Yunjie Zhao, Chen Zeng, Jianjin Xiang, Yi Zhang, and Yi Xiao. Optimization of RNA 3D structure prediction using evolutionary restraints of nucleotide-nucleotide interactions from direct coupling analysis. Nucleic acids research, 45(11):6299–6309, 2017.

32. Luqian Zheng, Christoph Falschlunger, Kaiyi Huang, Elisabeth Mairhofer, Shuguang Yuan, Juncheng Wang, Dinshaw J. Patel, Ronald Micura, and Aiming Ren. Hatchet ribozyme structure and implications for cleavage mechanism. Proceedings of the National Academy of Sciences, 116(22):10783–10791, 2019.

33. Jaswinder Singh, Jack Hanson, Kuldip Paliwal, and Yaoqi Zhou. RNA secondary structure prediction using an ensemble of two-dimensional deep neural networks and transfer learning. Nature Communications, 10(1):5407, 2019.

