## Supplementary material for "Improved RNA homology detection and alignment by automatic iterative search in an expanded database"

**Table S1:** Performance comparison among different DCA predictors on 245 PDB RNAs for RNAcmap2 and RNAcmap supplied alignment.

| | mfDCA | | | plmDCA | | | PLMC | | | GREMLIN | | |
|---|---|---|---|---|---|---|---|---|---|---|---|---|
| | F1 | Precision | Sensitivity | F1 | Precision | Sensitivity | F1 | Precision | Sensitivity | F1 | Precision | Sensitivity |
| RNAcmap | 0.436 | 0.494 | 0.397 | 0.425 | 0.482 | 0.387 | 0.417 | 0.474 | 0.378 | 0.404 | 0.460 | 0.365 |
| RNAcmap2 | 0.549 | 0.618 | 0.505 | 0.542 | 0.610 | 0.497 | 0.531 | 0.598 | 0.487 | 0.505 | 0.571 | 0.460 |

**Table S2:** Performance comparison among different MSA pipelines on No-hit RNAs (21 RNAs), Low $N_{eff}$ RNAs (83 RNAs), Medium $N_{eff}$ RNAs (31 RNAs), and High $N_{eff}$ RNAs (110 RNAs) using plmDCA predictor.

| | | No-hit RNAs | | | | Low $N_{eff}$ RNAs | | | | Medium $N_{eff}$ RNAs | | | | High $N_{eff}$ RNAs | | | |
|---|---|---|---|---|---|---|---|---|---|---|---|---|---|---|---|---|---|
| MSA | Pipeline | F1 | Precision | Sensitivity | Median $N_{eff}$ | F1 | Precision | Sensitivity | Median $N_{eff}$ | F1 | Precision | Sensitivity | Median $N_{eff}$ | F1 | Precision | Sensitivity | Median $N_{eff}$ |
| MSA-1 | BLAST-N | 0.000 | 0.000 | 0.000 | 0.0 | 0.010 | 0.011 | 0.010 | 1.0 | 0.024 | 0.028 | 0.021 | 1.0 | 0.045 | 0.052 | 0.040 | 2.1 |
| - | INFERNAL | 0.000 | 0.000 | 0.000 | 0.0 | 0.150 | 0.174 | 0.140 | 2.0 | 0.391 | 0.445 | 0.353 | 17.0 | 0.632 | 0.724 | 0.563 | 335.1 |
| MSA-2 | RNAcmap | 0.000 | 0.000 | 0.000 | 0.0 | 0.173 | 0.191 | 0.163 | 2.3 | 0.462 | 0.520 | 0.432 | 26.5 | 0.685 | 0.782 | 0.617 | 636.5 |
| MSA-2 | RNAcmap* | 0.125 | 0.133 | 0.128 | 1.0 | 0.225 | 0.249 | 0.211 | 4.1 | 0.511 | 0.577 | 0.475 | 31.6 | 0.697 | 0.795 | 0.628 | 605.1 |
| MSA-3 | RNAcmap2 | 0.159 | 0.171 | 0.159 | 1.4 | 0.416 | 0.457 | 0.389 | 13.0 | 0.589 | 0.661 | 0.542 | 123.2 | 0.697 | 0.796 | 0.629 | 605.1 |

**Table S3:** Performance comparison among different MSA pipelines on No-hit RNAs (21 RNAs), Low $N_{eff}$ RNAs (83 RNAs), Medium $N_{eff}$ RNAs (31 RNAs), and High $N_{eff}$ RNAs (110 RNAs) using PLMC predictor.

| | | No-hit RNAs | | | | Low $N_{eff}$ RNAs | | | | Medium $N_{eff}$ RNAs | | | | High $N_{eff}$ RNAs | | | |
|---|---|---|---|---|---|---|---|---|---|---|---|---|---|---|---|---|---|
| MSA | Pipeline | F1 | Precision | Sensitivity | Median $N_{eff}$ | F1 | Precision | Sensitivity | Median $N_{eff}$ | F1 | Precision | Sensitivity | Median $N_{eff}$ | F1 | Precision | Sensitivity | Median $N_{eff}$ |
| MSA-1 | BLAST-N | 0.000 | 0.000 | 0.000 | 0.0 | 0.015 | 0.017 | 0.014 | 1.0 | 0.015 | 0.018 | 0.013 | 1.0 | 0.006 | 0.007 | 0.005 | 2.1 |
| - | INFERNAL | 0.000 | 0.000 | 0.000 | 0.0 | 0.151 | 0.166 | 0.143 | 2.0 | 0.382 | 0.431 | 0.347 | 17.0 | 0.626 | 0.718 | 0.557 | 335.1 |
| MSA-2 | RNAcmap | 0.000 | 0.000 | 0.000 | 0.0 | 0.166 | 0.183 | 0.157 | 2.3 | 0.431 | 0.488 | 0.389 | 26.5 | 0.683 | 0.780 | 0.614 | 636.5 |
| MSA-2 | RNAcmap* | 0.131 | 0.146 | 0.122 | 1.0 | 0.216 | 0.238 | 0.203 | 4.1 | 0.482 | 0.544 | 0.450 | 31.6 | 0.690 | 0.788 | 0.623 | 605.1 |
| MSA-3 | RNAcmap2 | 0.158 | 0.173 | 0.156 | 1.4 | 0.404 | 0.443 | 0.379 | 13.0 | 0.557 | 0.625 | 0.514 | 123.2 | 0.690 | 0.788 | 0.623 | 605.1 |

**Table S4:** Performance comparison among different MSA pipelines on No-hit RNAs (21 RNAs), Low $N_{eff}$ RNAs (83 RNAs), Medium $N_{eff}$ RNAs (31 RNAs), and High $N_{eff}$ RNAs (110 RNAs) using GREMLIN predictor.

| | | No-hit RNAs | | | | Low $N_{eff}$ RNAs | | | | Medium $N_{eff}$ RNAs | | | | High $N_{eff}$ RNAs | | | |
|---|---|---|---|---|---|---|---|---|---|---|---|---|---|---|---|---|---|
| MSA | Pipeline | F1 | Precision | Sensitivity | Median $N_{eff}$ | F1 | Precision | Sensitivity | Median $N_{eff}$ | F1 | Precision | Sensitivity | Median $N_{eff}$ | F1 | Precision | Sensitivity | Median $N_{eff}$ |
| MSA-1 | BLAST-N | 0.000 | 0.000 | 0.000 | 0.0 | 0.006 | 0.007 | 0.007 | 1.0 | 0.035 | 0.040 | 0.032 | 1.0 | 0.074 | 0.087 | 0.066 | 2.1 |
| - | INFERNAL | 0.000 | 0.000 | 0.000 | 0.0 | 0.150 | 0.163 | 0.141 | 2.0 | 0.343 | 0.395 | 0.307 | 17.0 | 0.607 | 0.698 | 0.540 | 335.1 |
| MSA-2 | RNAcmap | 0.000 | 0.000 | 0.000 | 0.0 | 0.165 | 0.180 | 0.156 | 2.3 | 0.404 | 0.460 | 0.368 | 26.5 | 0.663 | 0.760 | 0.591 | 636.5 |
| MSA-2 | RNAcmap* | 0.115 | 0.121 | 0.118 | 1.0 | 0.213 | 0.231 | 0.204 | 4.1 | 0.457 | 0.518 | 0.426 | 31.6 | 0.663 | 0.761 | 0.591 | 605.1 |
| MSA-3 | RNAcmap2 | 0.133 | 0.145 | 0.137 | 1.4 | 0.380 | 0.418 | 0.357 | 13.0 | 0.531 | 0.598 | 0.487 | 123.2 | 0.663 | 0.761 | 0.591 | 605.1 |

**Table S5:** Performance comparison of RNAcmap* and RNAcmap2 supplied alignments for RNAfold and SPOT-RNA as Consensus Secondary Structure (CSS) using mfDCA predictor on 102 PDB RNAs non-redundant from SPOT-RNA training data at 0.8 sequence identity cut-off using CD-HIT-EST.

| | F1 | Precision | Sensitivity | Median $N_{eff}$ |
|---|---|---|---|---|
| RNAcmap* (RNAfold) | 0.398 | 0.438 | 0.377 | 12.2 |
| RNAcmap* (SPOT-RNA) | 0.434 | 0.472 | 0.425 | 19.5 |
| RNAcmap2 (RNAfold) | 0.499 | 0.545 | 0.476 | 55.6 |
| RNAcmap2 (SPOT-RNA) | 0.516 | 0.559 | 0.503 | 57.6 |

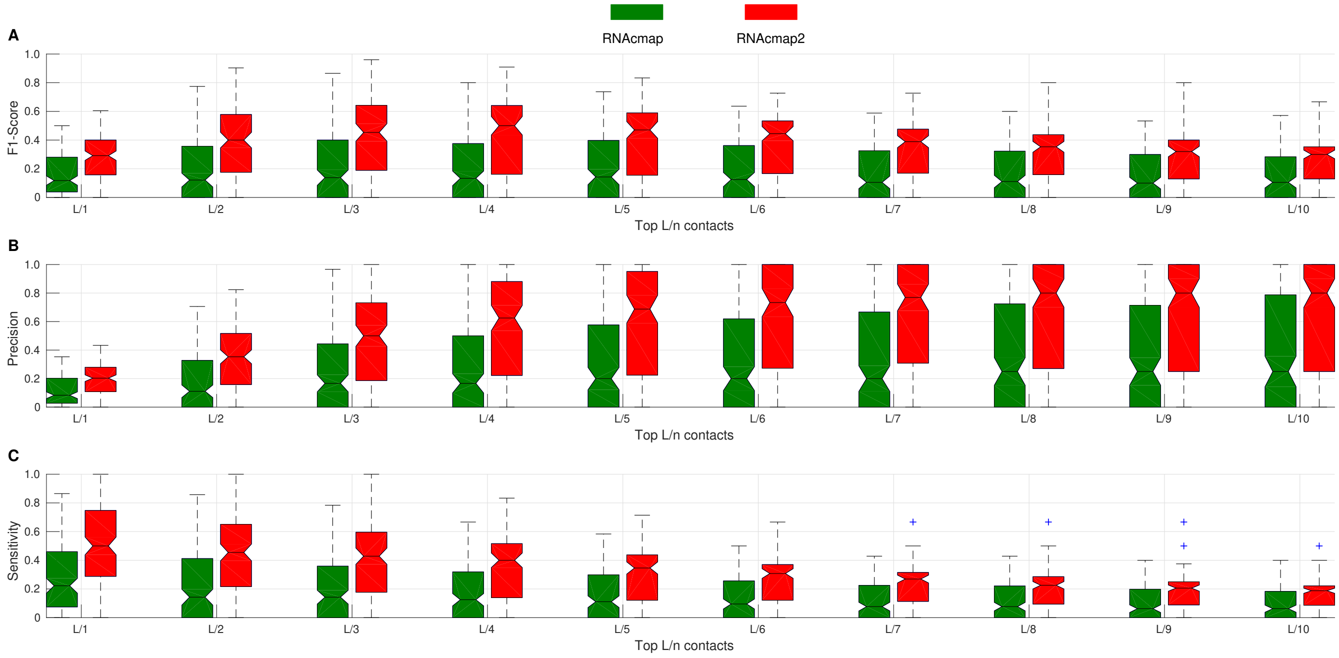


**Figure S1.** Boxplot of F1-score (**A**), Precision (**B**), and Sensitivity (**C**) as a function of predicted top L/n base pairs by plmDCA from RNAcmap (in green) and RNAcmap2 (in red) supplied alignment for 135 PDB RNAs from no-hit, low, medium $N_{eff}$ test sets. The distribution is shown in terms of median, 25th and 75th percentile with outlier shown by dots.

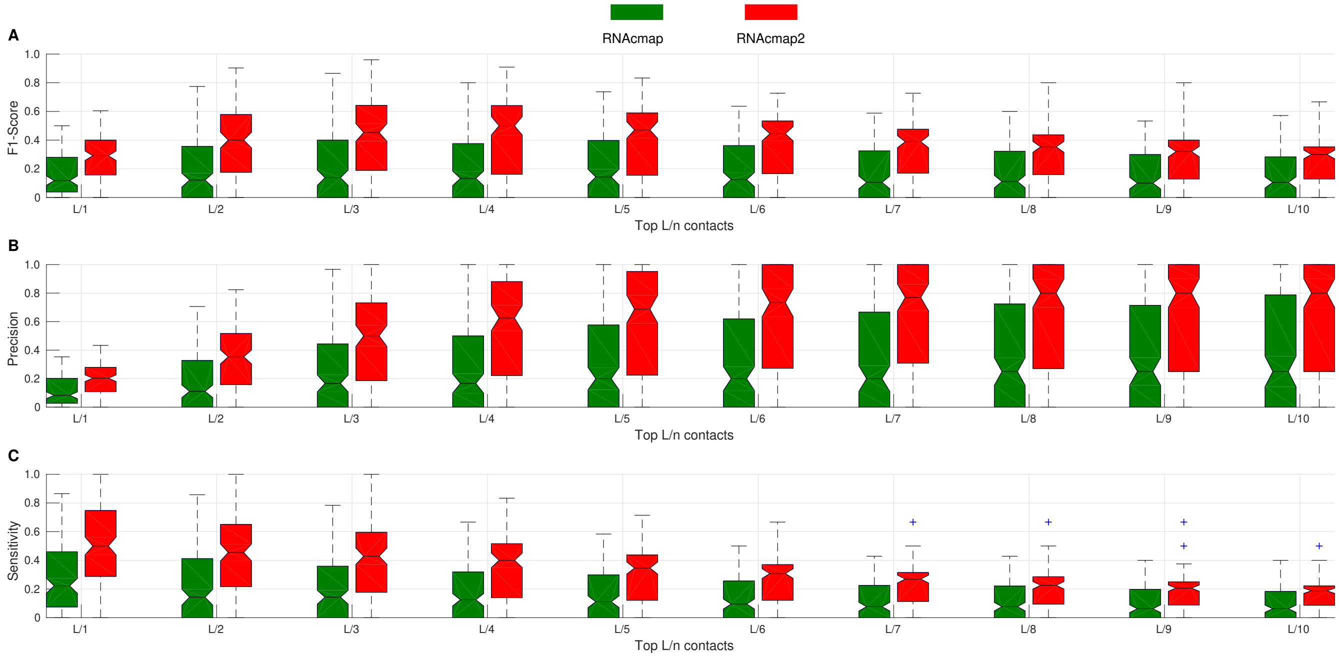


**Figure S2.** Boxplot of F1-score (**A**), Precision (**B**), and Sensitivity (**C**) as a function of predicted top L/n base pairs by PLMC from RNAcmap (in green) and RNAcmap2 (in red) supplied alignment for 135 PDB RNAs from no-hit, low, medium $N_{eff}$ test sets. The distribution is shown in terms of median, 25th and 75th percentile with outlier shown by dots.

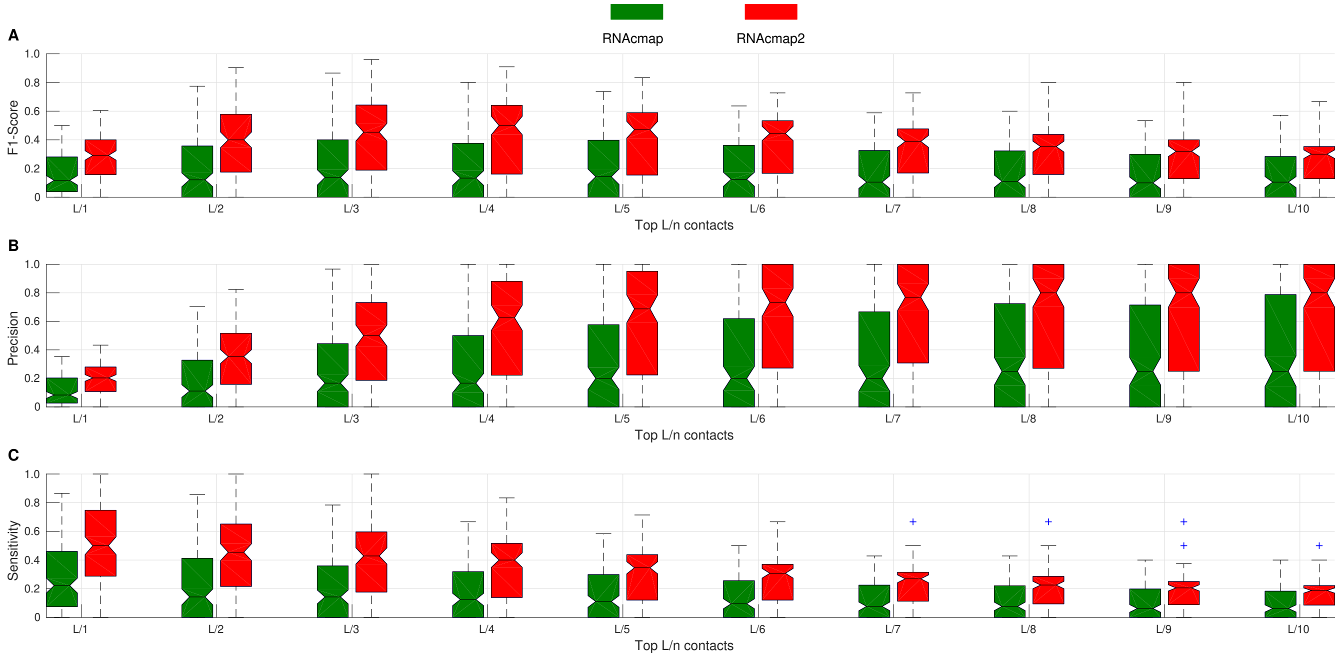


**Figure S3.** Boxplot of F1-score (**A**), Precision (**B**), and Sensitivity (**C**) as a function of predicted top L/n base pairs by GREMLIN from RNAcmap (in green) and RNAcmap2 (in red) supplied alignment for 135 PDB RNAs from no-hit, low, medium $N_{eff}$ test sets. The distribution is shown in terms of median, 25th and 75th percentile with outlier shown by dots.
